# OrganL: Dynamic Triangulation of Biomembranes using Curved Elements

**DOI:** 10.1101/2023.09.18.557233

**Authors:** Christoph Allolio, Balázs Fábián, Mark Dostalík

## Abstract

We describe a method for simulating biomembranes of arbitrary shape. In contrast to other dynamically triangulated surface (DTS) algorithms, our method provides a rich, *quasi* tangent-continuous, yet local description of the surface. We use curved Nagata triangles, which we generalize to cubic order to achieve the requisite flexibility. The resulting interpolation can be constructed locally without iterations. This allows us to provide a parallelized and fine-tuned Monte Carlo implementation. As a first example of the potential benefits of the enhanced description, our method supports inhomogeneous lipid properties as well as lipid mixing. It also supports restraints and constraints of various types and is constructed to be as easily extensible as possible. We validate the approach by testing its numerical accuracy, followed by reproducing the known Helfrich solutions for shapes with rotational symmetry. Finally, we present some example applications, including curvature-driven demixing and stylized effects of proteins. Input files for these examples, as well as the implementation itself, are freely available for researchers under the name OrganL.

Our method provides a straightforward way to simulate any biomembrane geometry. It overcomes some of the limitations of previous dynamically triangulated surface (DTS) Monte Carlo schemes by providing a surface that contains an interpolant which allows to assign meaningful functions of curvature to almost every point of the discretization, yet keeps much of the simplicity of the common DTS schemes by not requiring any nonlocal information or iterations for its construction. Our tool is easily extensible and facilitates the simulation of complex lipid and protein compositions on membrane surfaces at any scale.

## INTRODUCTION

A realistic multiscale model of (sub)cellular structures, such as organelles remains a distant goal. Molecular dynamics (MD) simulations provide detailed information on molecular structure and surface binding but are limited by size and time scales. Coarse-grained (CG) MD simulations provide limited relief, as they still require to simulate particles of a definite size and often entail a substantial loss of accuracy and transferability.(1) The obvious starting point for the mesoscopic description of lipid membrane-based systems is provided by the (Evans-Canham)-Helfrich functional(2–4), which has found wide application in membrane biophysics(5–9) since its introduction in the seventies. Recently, efficient techniques to extract its local parameters from atomistic MD simulations have been developed.(10–14) Introducing molecular specificity into the continuum model in this way has the potential to allow the systematic construction of models for slow processes at the cellular scale and to solve them using modest computational means. An illustration was recently provided for membrane fusion.(11) The heterogeneous nature of biomembranes requires us to provide local and dynamic membrane properties. However, even in the absence of such concerns, the Helfrich functional is difficult to extremalize analytically as well as numerically. This is particularly true in the absence of symmetries, as surfaces are hard to parameterize in a general setting. DTS simulations, as pioneered by Kroll and Gompper(15) among others(16–18) are, therefore, often employed to explore static as well as time-dependent behavior of complex membrane structures.(19) The DTS method, as it is established, is based on an operator discretization at the vertex.(20) Such discretizations are widely used in known packages(21, 22), such as Brakke’s surface evolver(23), but the underlying construction does not allow for any substructure on the faces, relegating all information to the vertices. This leads, for example, to the practice of confining membrane-deforming proteins to the mesh-vertices(22, 24) and makes it difficult to compute accurate interaction energies of membrane structures at curved interfaces. The insertion of complex mixing and coupling terms or flow fields thus requires a very fine mesh, even though actual changes in shape are often of large scale when compared to the protein size. It is, of course, possible to describe curvature in a continuous manner, using splines(25) and there are other known finite element approaches.(26) The Nagata interpolation(27) is *quasi* tangent continuous and, as we shall see, provides a sufficient degree of accuracy in the calculation of the curvature integrals required for the Helfrich theory. However, it also has substantial drawbacks. In particular, the interpolation requires the provision of surface normals. Normal estimation on unstructured meshes remains a challenging problem and the standard Nagata interpolant fails to give reasonable results for a wide range of “unexpected” normals.(28) As the resolution of the problems of the Nagata interpolant requires modification of both the interpolant and commonly used Monte Carlo steps, the Methods section necessarily contains a significant amount of new methodology; in other words, it contains results.

## METHODS

### Discretization Scheme

The starting point of the Nagata interpolation scheme(27) is the search for a quadratic edge interpolant ***φ*** (t) between two vertices ***x*** _*A*_, ***x***_*B*_. The distance vector ***d*** = ***x***_*B*−_ ***x*** _*A*_ connects the two vertices. We will denote the coefficients (vector) of the interpolant with *c*_1_. Defining

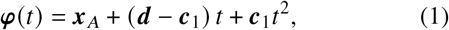

ensures that the interpolant goes to ***x*** _*A*_ at *t* = 0 and ***x***_*B*_ at *t* = 1. Furthermore, if the coefficient vector ***c***_1_ is zero, ***φ*** (t) becomes a straight line connecting the vertices. The key difference to other interpolation schemes is that instead of solving for the coefficients by some boundary conditions (such as equal derivative values) defined by the neighboring patches, the Nagata scheme uses normal vectors ***n***_1_, ***n***_2_ to determine the coefficients without any direct reference to neighboring patches. Reproducing surface normal vectors requires the knowledge of at least one more interpolated edge to construct the tangent basis. However, for simplicity, Nagata decided to merely consider tangent vectors of the single edge interpolant

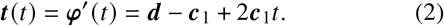

A consistent normal vector requires the tangents ***t*** _*A*_ = ***t*** (0) and ***t***_*B*_ = ***t*** (1) at the endpoints to be orthogonal to the normal.Hence, the conditions

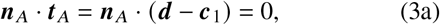

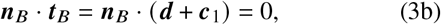

need to be fulfilled. However, ***c***_1_ has three elements. Hence the resulting linear equation system 𝕄 for ***c***_1_ is underdetermined. As ***c***_1_ controls the “curvature” of the interpolant, it makes sense to insert the minimization of ||***c***_1_|| as the third condition to complete the system. Note that for || ***c***_1_|| > 0, this norm has no straightforward relation with curvature as it is commonly defined on curves. The pseudoinverse, also known as the Moore-Penrose inverse of a matrix 𝕄, is a generalization of the matrix inverse to nonsquare matrices. It guarantees fulfillment of 𝕄, while minimizing ||***c***_1_|| (least squares). Following reference (27), the pseudoinverse can be computed analytically via

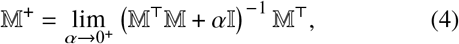

leading to simple expressions for the determination of ***c***_1_. Fig. 1 shows the construction of an edge interpolant in a non-pathological case.

**Figure 1.**
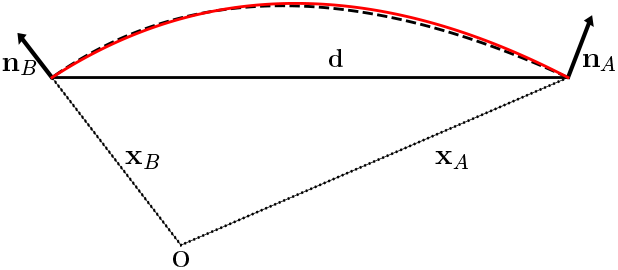
Quadratic (dotted line) and cubic (red line) Nagata edge interpolants, for the case of well-behaved normal vectors.

Unfortunately, this process fails to give the expected results for a wide variety of pathological normal orientations. Singular points and their remedies have been discussed by Nagata.(27) For our purposes, it is sufficient that the interpolation will succeed for smooth surfaces, as is required by the ordinary Helfrich theory. It was discovered recently that the results for

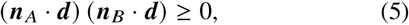

are problematic.(28) Some of the issues are illustrated in Fig. 2. Namely, if both of the normal vectors are oriented in the same way with respect to ***d***, the quadratic interpolant will not lie in the angle between normal vectors viewed along ***d***. An example of this arrangement is given in the left panel of Fig. 2. Namely, the two normals are approximately parallel and close to orthogonal to ***d***. Such a configuration of normal vectors may arise readily, e.g. from a normal estimation with a small perturbation of a flat mesh during the formation of a dimple. The resulting interpolant describes a large “sideways” bow, risking overlaps with other parts of the mesh. Similar problems occur in the event of twisting the normals as shown on the right of Fig. 2. If the interpolant were to lie within the space spanned by the normals and ***d***, it would have to contain an inflection point. For such cases, Nagata and subsequent authors advise either subdivide the mesh to cover the inflection point or to insert a flat line. In the case of a membrane simulation, such an approach is difficult to implement, as there are no underlying data to refine and a realistic expression of curvature is required. In order to ameliorate the situation, we increase the interpolation to cubic order:

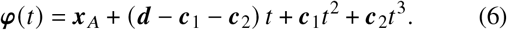

The derivative is given by

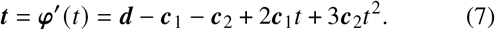

As before, we impose the requirement of the tangent vectors to be perpendicular to the normals at the endpoints. The introduction of ***c***_2_ means that the equation system now has four degrees of freedom left to determine. Note that, e.g. the arc-length variation of a curve between two points, for a given set of tangent vectors will result in a straight line with an infinitesimal kink; it is therefore expected that an arbitrary increase of polynomial order will lead to an equivalent result. In order to gain a good interpolant, it is necessary to require additional conditions. For the pseudoinverse operation to work, these conditions have to come in the form of linear equations. Therefore, we impose an additional estimate of the binormal vector. This vector is orthogonal to the normal vector and is estimated to be normal to the plane spanned by ***n*** and ***d***. This leads to the following set of conditions:

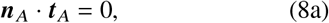

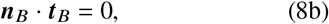

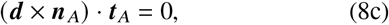

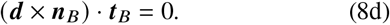

As remarked previously, using the pseudoinverse for an underdetermined system corresponds to solving a least squares problem in the coefficient vector. This problem has an analytical solution, which we provide in the Supplementary Material, together with the detailed algorithm which produces the edge interpolant with a few vector multiplications. We find them to be well-behaved, resulting in an interpolant very close to the original Nagata formulation for the “nonpathological” cases as well as a reasonable inflection point for the failure mode described in Eq. (5). The additional condition introduced minimizes the chances of edge overlap.

**Figure 2.**
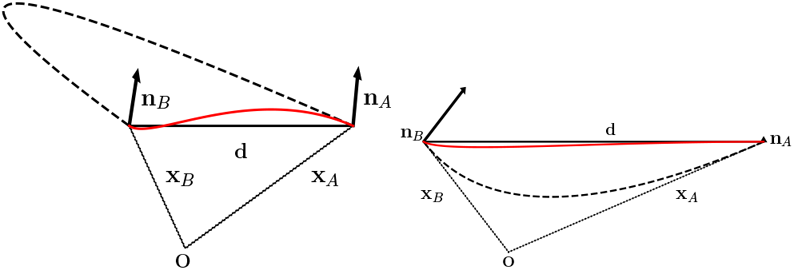
Left panel: Quadratic (dotted line) and cubic (red line) Nagata edge interpolants, for the case of normal vectors with equal orientation ***n***_*i*_ · ***d*** > 0. Right panel: Arrangement for twisted normals with the additional condition of ***n***_2_ · ***d*** ≈ 0.

### Interpolation of a Patch

In order to create a surface interpolant, the edge interpolants have to be assembled into a surface. The patch interpolant ***x***_2_(*η, ζ*) for the original quadratic Nagata edges is also of second order:

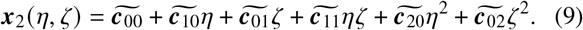

The six coefficients of this polynomial are determined by the requirement of the edges of the surface interpolant being equal to the edge interpolants. More precisely, the functions ***x***_2_ (*η*, 0), ***x***_2_ (1, *ζ*), ***x***_2_ (*η, η*) have to be exactly equal to the Nagata edge interpolants. For a visualization of the structure, see Fig. 3. The edge polynomials restrict the domain of the interpolated patch to 0 ≤ *η* ≤ 1 and 0 ≤ *ζ* ≤ *η*.

**Figure 3.**
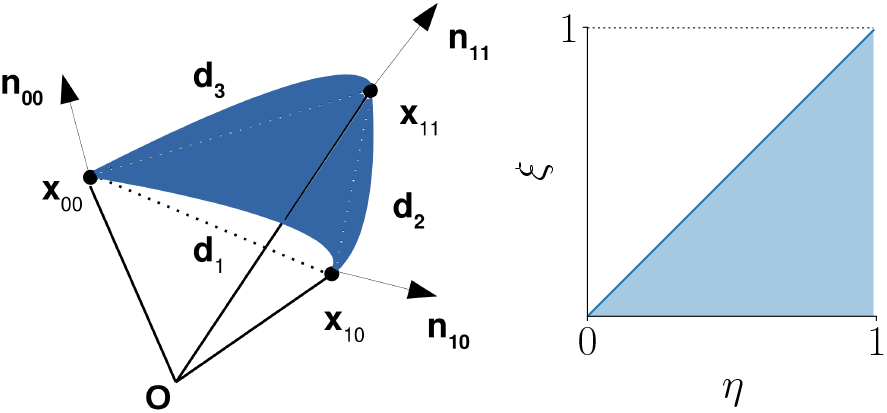
Left panel: Schematic of a patch interpolant as constructed from the edge interpolants. Right panel: Domain of the patch interpolant.

The third-order patch interpolant requires additional conditions to be uniquely determined. We require it to become identical to the quadratic case in case of ***c***_2_ = 0. This resolves the ambiguity. A detailed derivation is given in the Supplementary Material. In our code, cubic edges are only used for those edges, where the failure condition in Eq. (5) is met. The alternatives of permitting bad interpolants or inserting a straight line lead to heavy artifacts. Excluding these configurations leads to the necessity of creating complicated, collective motions of the mesh (clustered moves) to avoid trapping. The stand-alone accuracy of the resulting interpolant is evaluated in the Results section.

### Evaluation of the Helfrich Energy

The Helfrich free energy F, supplemented by surface tension σ and pressure *p* terms can be stated as

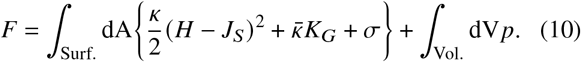

In this arrangement, the geometrical quantities include the mean curvature *H*, the Gaussian curvature *k*_*G*_, the area element dA and the volume element dV. The material parameters are the bending rigidity *κ*, the Gaussian bending modulus 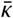 and the spontaneous or intrinsic curvature *J*_*S*_. The pressure *p* and surface tension σ can be interpreted as Lagrange multipliers on the total surface and volume, respectively. In our scheme, *A* and *V* are controlled instead by quadratic penalties and *k*_*G*_ is ignored for topological reasons. We note, that *V* can be obtained from the divergence theorem using the spatial coordinate ***r*** as

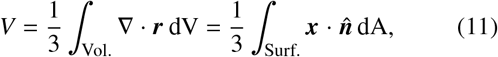

where 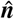 is the surface unit normal vector as computed from the patch interpolant ***x***. Therefore, it is sufficient to work with surface integrals to evaluate all of F. The surface integral is summed over all patches and in each patch is discretized on the domain of the interpolant using a seventh-order Gaussian quadrature scheme(29). Note the facility of, e.g. constructing parallel planes accounting explicitly for separate monolayers and the availability of a reasonable direct way to evaluate Gaussian curvature on a face. On the level of the code, different energy functionals are available via subclassing. Some are described in the Supplementary Material, most notably an implementation of area difference elasticity (ADE)(30, 31) via control of the integrated mean curvature. Local curvatures can be expressed in a simplified manner by the coefficients. More details about the implementation are given in the Supplementary Material.

### Parallelized MC Implementation

#### Moves

We use a standard Metropolis Monte-Carlo (MC) algorithm to evolve the interpolation and sample the Boltzmann distribution of the underlying energy. In contrast to the standard DTS, we also have to consider the changes of the normals in the sampling of equilibrium geometries. We use vertex moves, normal moves and so-called deep vertex moves. In addition to this, we have also implemented a basic remeshing “Alexander” move and a new type of move associated with lipid composition on a face. Due to the local nature of these moves, the code is easily parallelizable. In Fig. 4, we show the part of the local mesh that is modified, as it needs to be checked for validity and is updated in every step. The **normal move** does not change any of the mesh vertices. In order to generate the move, we create a vector ***d*** from a Gaussian distribution

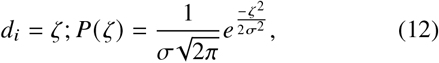

where the components *d*_*i*_ are set to separate Gaussian random variables. The pseudorandom variables *ζ* are generated using a Mersenne-Twister algorithm, initialized by a time-dependent seed. The proposed new normal vector is then obtained as

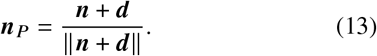

Note that the probability of proposing −***d*** is the same as the probability of proposing ***d***, due to the symmetry of the Gaussian distribution. σ can be adjusted automatically. For proposal probability, we find

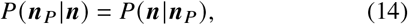

which helps to ensure detailed balance and allows us to compute the acceptance probability as

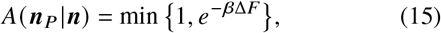

where ΔF is the change in the total free energy associated with the transition and β = (*kT*) ^−1^. We state F in units of *kT*, cancelling β. The change in energy can be computed locally by considering only the triangles which contain this normal, hence it is trivially parallelizable. The only difficulty pertains to the evaluation of changes to the global quantities *V* and *A*. For a global “restraint” on a function Φ to Φ_0_, with an energy contribution of the form

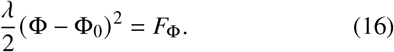

The change ΔΦ due to a local MC step leads to an energy change of

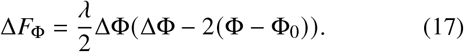

This means, that a local scheme requires the storage of the current global value Φ. However, Φ can be updated using ΔΦ if the current local contribution to Φ is stored on the face. In order to accept the proposed normal, it will also have to produce valid cubic or quadratic face interpolants. To ensure this, we require the proposed normal to be at a minimum angle to the other triangle normals ***n***_*i*_, so that

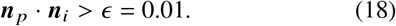

Finally, a collision check is performed. This collision check is delayed because of the great computation expense of evaluating the neighbor list. The second move to consider is the **vertex move**. Again, a vector is sampled from a normal distribution and added to the previous position. The same sort of symmetry argument, leading to Eq. (14) applies. The change of a vertex has more spatially extended consequences than the change of a normal, see Fig. 4, b). Of course, the triangles in the immediate vicinity of the initial vertex have to be updated. The acceptance criterion in Eq. (15) is applied. Moreover, the acceptance of the vertex move requires

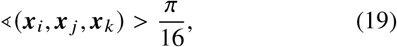

for all of the angles in the triangle. The “**deep-vertex**” move from Fig. 4c, assumes that the normal vectors are not independent degrees of freedom, but are determined by the faces of the underlying triangular mesh. Under these conditions, the move is fully reversible, as it becomes equivalent to a vertex move with automatic restoration of the normal estimate. This would not necessarily be true if the move was to only update the normal on its own vertex, because under these conditions the vertex normals of the surrounding faces would no longer correspond to their estimated values. As a remedy, we recommend to “quench” the normals to their estimated values before using this move or to use it standalone. Our MC scheme is also set-up in such a way as to minimize irreversibility by the organization of the Markov chain. In particular, for each degree of freedom, multiple trials(32) are proposed until acceptance, so that for each MC step (almost) all degrees of freedom have moved. The total step can therefore be interpreted as a factorization of individual symmetric steps.(33) Furthermore, we have implemented a mechanism of automated propagation, which ensures all other energy carrying objects are updated together with the faces. Algorithmic details on the MC scheme and the automatic construction of decoupled parallel execution loads are given in the Supplementary Material. The **Alexander move** d) is currently set to be accepted only if it decreases energy. Our automatic parallelization scheme, described in the SI relies on tracking the dependencies of each degree of freedom as determined from mesh. As the Alexander move changes the mesh topology, the parallelization structure of the code needs to be updated, and neighbor lists rebuilt after each round of Alexander moves. Hence, the remeshing move is executed only occasionally and (mostly) serially. As the number of faces is conserved, face properties (such as local bending rigidity) can remain associated with a fixed face, currently, no redistribution occurs. The Alexander move is associated with the edge which is flipped. The **Lipid mixing** move e) is also edge-associated, random lipids are chosen from the population of both faces and transferred to another face. More information is found in the lipid mixing section.

**Figure 4.**
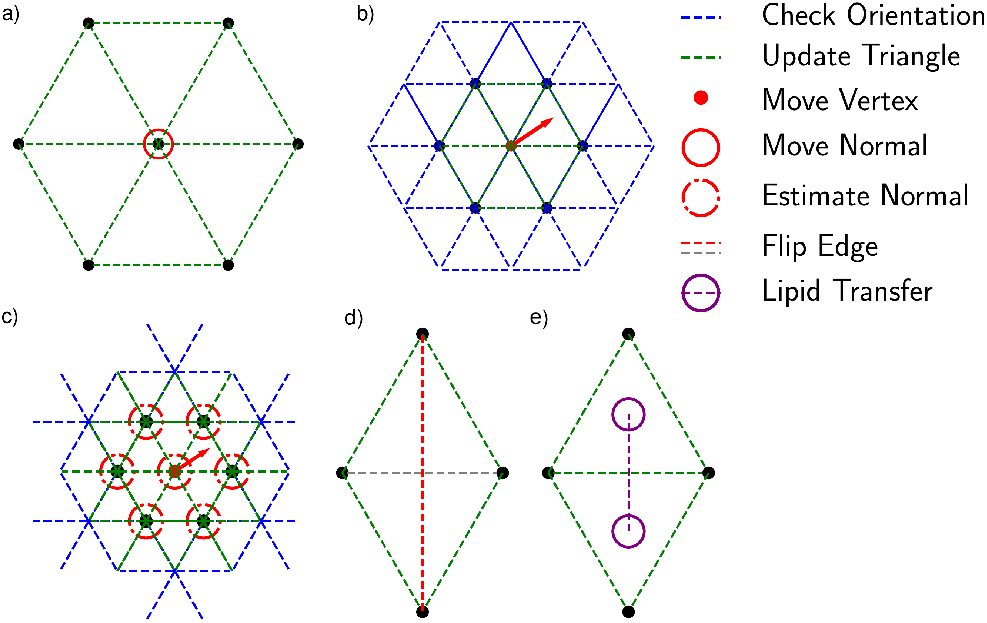
Implemented MC-Moves: a) Normal move and its update range, b) vertex move and it is update and face orientation check range, c) Deep vertex move, d) Alexander move, e) Lipid transfer move. The legend describes the update schemes, face updates and orientation checks.

### Mesh Properties and Constraints

#### Constraints

It is possible to selectively block or remove parts of the mesh from active evolution. The former can be accomplished by removing some properties while retaining the mesh points.

The latter is accomplished via *Block* objects, which suppress updating of parts e.g. only the *x* coordinate of a vertex or normal. Thus, a simple way of implementing sophisticated boundary conditions is available. An automatic way to load in simple coordinate and normal freezes via text files is provided.

#### Face Properties

As energy and integral evaluations are performed over the faces, it makes sense to also store information about the parameters *κ, J*_*s*_, *A*_0_ and others on them. In fact, each face contains an arbitrary collection of properties, which are accessed via a text key. Such properties can be assigned at load as well as at run time and are automatically included in output visualizations. This allows us to dynamically store and export information about composition as well as compute modified elastic properties and local coupling terms on the fly.

### Lipid Mixing

The plethora of easily implemented energy functionals would also allow to compute the energy directly from a composition. However, a proper implementation of lipid mixing requires the lipid composition to be dynamic and, therefore, requires the definition of mixing rules and moves which manipulate it. Ideally, the modification of the lipid composition should be transparent to most of the energies computed from it. In the literature, it has often been proposed to compute the bending rigidity *κ* as(34, 35) via a harmonic average of the single lipid moduli *k*_*i*_:

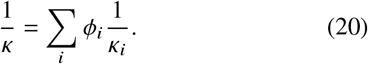

Here *ø*_*i*_ refers to the volume fraction, which is calculated as

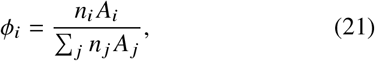

with *A*_*i*_ the area per lipid of the lipid species *i*. This means, that we assume that the area per lipid does not change while mixing. By *n*_*i*_, we refer to the number of lipids of species *i* present in a triangle. Of course, completely different behavior is possible. An example occurs during a phase transition to gel or L_*O*_ phase. As long as the behavior is well-defined it is easy to implement the requisite models in our code. The spontaneous curvature is computed as

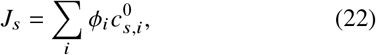

where 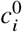 is spontaneous curvature of the pure *i*.(13, 36) We have recently tested the validity of these equations against atomistic molecular dynamics results.(37) Furthermore, the mixing entropy of the lipid composition can be estimated as ideal in the surface fractions(36):

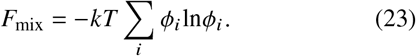

This free energy is added to the energy in the form of a penalty. In this way, it can be used with all energy functionals. It is worth pointing out that real valued molar fractions *φ*_*i*_ and lipid occupancies *n*_*i*_ presume that the lipid composition of a triangle is a macroscopic quantity. As the mixing entropy is temperature dependent, the temperature is provided via the inverse temperature β, which can be set in a simulation-wide manner. Furthermore, both leaflets in a bilayer are considered, and their properties are computed separately, and then added together. The equilibrium area of a face *A*_0_ is computed from the areas per lipid on each leaflet, which are then averaged. There is also the possibility to use a stylized protein, whose move will be integer valued and which will override the curvature generated by the lipids. In order to simulate lipid mixing, an exchange move has been implemented. This move works by the following scheme. On each leaflet:

1. Select two lipid types at random from adjacent faces.
2. Generate two random amounts of lipids to be moved.
3. Move the two lipids between the adjacent faces and recompute the energies, including mixing energy.
4. Accept or reject the move.

Uniform random numbers are used for the exchange move. In order for the move to be allowed, the total occupancy must be above a certain threshold, e.g. zero. Currently, the moves are set so that the total area of each face and leaflet does not change by lipid exchange. This serves to prevent mesh degeneration.

### Collision Detection

The detection of potential collisions between any pair of elements in a set of *n* triangles requires O (*n*^2^) operations. To mitigate the quadratic time complexity of the collision detection, we implemented a neighbor-list based *broad* (38) search that aims to reduce the number of explicit intersections which have to be computed. The broad search is followed by a *narrow* phase that relies on the popular (39–41) fast triangle-triangle intersection test of Möller (42). As the mesh faces are cubic surfaces, the use of the Möller algorithm is a compromise between precision and computational cost.

### Penalties and Errors

It is difficult to quantify the discretization error on a Nagata patch. However, we can require that its integral is consistently defined across the entire surface. This is not necessarily the case for the Nagata interpolant. At a patch boundary one of the tangent vectors, i.e. the one taken along the edge interpolant is identical to that of its neighboring patch. The other tangent vector may be different so that for an identical point at the edge two different tangent vectors are obtained. For the calculation of the energy, and even the consistency of the interpolation, we require a consistent definition of the surface integral everywhere on the surface and to evaluate the same for an arbitrary subdomain

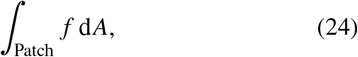

where *f* may depend on derivatives of the surface or might be a differential form. For example, a smooth surface in R^3^ will fulfil

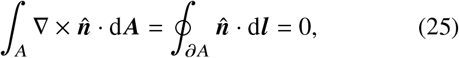

for any closed, finite area *A* with boundary *∂ A*, tangent vector d***l*** and oriented surface element d*A*, and. It is easily shown that 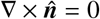 is fulfilled everywhere inside an interpolated patch.If d*l* is a tangent vector of an edge interpolant, integrating around the boundary of each patch will also fulfill the condition as even where normals are ambiguous/discontinuous on the edge interpolant, both are orthogonal to the tangent vector which follows the boundary line by construction. However, we need more regularity than this, as we are interested in integrals of *H* and *H*^2^; in general, our interpolant is only C^0^ continuous, across edges but the computation of curvature requires derivatives of ***n*** in any direction, in particular the mean curvature *H*, i.e. the covariant normal divergence

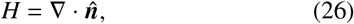

has to be (square) integrable on the surface. At the edge, one tangent vector is shared (hence the same normal curvature in one direction(43)). If the normal vector at this point is also identical, it is possible to construct a local orthogonal tangent basis, this second tangent defines 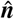 as it stands orthogonal to the edge direction and ***n*** at all times. Tangent, i.e. C ^1^ continuity, intuitively is sufficient to guarantee some curvature integral properties, but does not guarantee continuity of the curvature itself. It came as a surprise to us, that the patch interpolant does not always reproduce the vertex normals used to create it, but closer examination shows that this is not guaranteed, as e.g. one of the edge interpolants during construction may have parallel tangents at the endpoints or might generate normal vectors of opposite orientation. Taking a pragmatic approach, we will simply make sure that

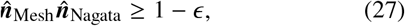

before accepting a move; we use *ϵ* = 0.005. In order to bring the discretization closer to a C^1^ continuity, we first define the following energy scale

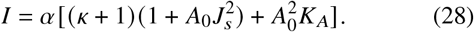

The individual terms in this intensity are spatial-scale free energy units. They are set to cover all sources of energy in the problem so that a significant *J*_*s*_ does not damage the regularizing penalty. Now, we split the tangent basis into two parts,

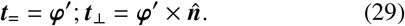

We then require that at the midpoint of an edge

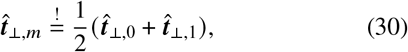

where the subscripts (0,1) denote the normalized average value of ***t*** at the endpoints of the edge interpolants. By controlling the evolution of 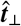, in each triangle, we enforce the continuity of normals at the center of the triangle, biasing towards C^1^ continuity without requiring information from adjacent triangles. Deviation from Eq. 30 carries a quadratic penalty depending on *I*. We also penalize too small triangle areas (starting < 40%*A*_0_) and deviations from an equilibrium angle (| *α*| < 0.316) with quadratic penalties proportional to *I*. The interpolant is highly tolerant of different penalties, as long as they are sufficiently strong. The area penalty serves mainly to prevent mesh degeneration. This mesh degeneration is less of a problem for the interpolant than for the MC algorithm, as the admissible step size becomes highly in-homogeneous. Similarly, the angle penalty mostly serves to prevent the locking and overlap of triangles.

### Input and Output Formats

The mesh input file format for this code is wavefront obj. This format can be exported from most free and standard tools. If the original obj file does not contain normals, they will be reconstructed from neighboring face normals in a simple weighting scheme(44). The program also exports this type of mesh, in addition to XML-VTK(45) unstructured grid files (.vtu). These are readily available and can be visualized and processed with the Paraview software package(46).

## RESULTS AND DISCUSSION

### Example Meshes

It is difficult to unit-test the error of the interpolation. The easiest way to proceed is to demonstrate the accuracy for example meshes, using reconstructed normals. The results are shown in Fig. 5. The irregular mesh for the catenoid, combined with the difficult normal estimation at the edges, results in a total curvature integral of 0.8 overall faces and a total Helfrich energy of 0.6*κ* over an area of 183.585 units. Any minimal surface has *H* = 0 everywhere, so the theoretical value for mean curvature integrals is zero. For the unit icosphere mesh, the corresponding values are a total curvature of 12.5662, energy of 12.4780*κ* and area of 12.5662. The theoretical values for energy, area, curvature are 4*π*(*κ*) ≈12.5664 as *R* = 1. The error for volume is negligible as well. This illustrates the high accuracy of our scheme for good quality meshes.

**Figure 5.**
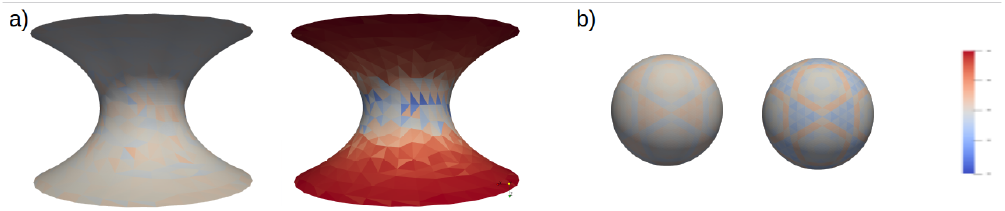
Two example meshes a) a medium quality mesh of a catenoid geometry (1096 faces, 580 vertices), b) Icosphere mesh (642 vertices, 1280 faces) with mean curvatures and gaussian curvatures plotted as averages over face integrals. The scales are adjusted to run from the theoretical value ± 20 % of the maximum value for the smallest radius of curvature.

We also tested the icosphere using purely cubic interpolants. Area and volume accuracy remained unaffected, but energy decreased to 12.340*κ* and curvature to 12.44. For the catenoid, energies increased to 0.85*κ*, while volume and area remained indistinguishable. We note, that the quadratic interpolation appears more robust, but errors remain small. At present, we do not know how important integration order is for error control. We fall back on the quadratic interpolant, when normals allow it.

### Helfrich Minima

In order to validate the interpolant and our MC algorithms, we reproduced the behavior of the known rotationally symmetric solutions of the Helfrich Hamiltonian. The minima have been well explored and confirmed recently by independent techniques.(20, 25) Results are shown in Fig. 6. We have generated all the points using a discocyte starting geometry. In particular, thanks to the MC implementation we are able to transform prolate to discocyte and vice-versa near the theoretical limits, which are indicated by the dotted line. The prolate state can be somewhat metastable, so that, with fast pulling some “wormlike” shapes were obtained. The transition from discocyte to stomatocyte was also observed and occurred via collision and repulsion of the stomatocyte dimples. The theoretical curve is shifted by the thermal energy *α*. The value of alpha was estimated by heating the spherical geometry and by annealing to be ≈ 0.7. The geometries shown in Fig. 6 also reveal the good quality of the curvature on each individual face (only the average values are mapped on each face).

**Figure 6.**
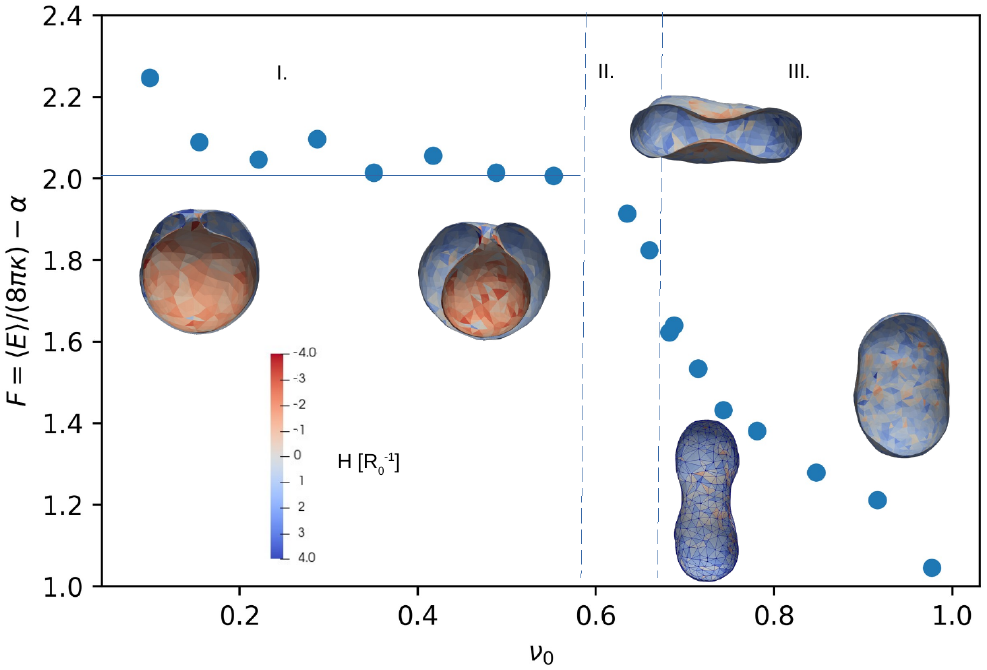
Results for spherical vesicles, including the minima for stomatocytes I., discocytes II., discoid and oblate III. shapes obtained using volume and area constraints. The blue line indicates the theoretical value for the stomatocyte. *α* is due to the thermal energy of the degrees of freedom of the mesh.

For this simulation, only the global surface area *A* and the global volume *V* were restrained. From these was computed the reduced volume

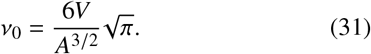

When the space inside the vesicle becomes too small, i.e. at *v*_0_ < 0.2 fluctuations deform the mesh and the neck becomes heavily penalized. While we believe it might be possible to go even lower with appropriate penalties, this seems to be quite sufficient for most stomatocyte shapes.

### Examples: ADE and Lipid Mixing

The sampling of the standard Helfrich minima in Fig. 6 is just a precondition for the more advanced methods. With a robust description of the surface, it also becomes possible to add the lipid moves described in the previous section. For the purposes of clarity, we have chosen only one leaflet to have a strong curvature. In Fig. 7 a), the comparison of a mixed patch with *J*_*s*_ = 0 is shown. The solver spontaneously generates a protrusion and lipid demixing driven by curvature. The lipid population and local *J*_*s*_ are visualized together in central and right panels. Note, how the boundary of the simulation is constrained. The starting geometry was a mesh-enhanced spherical cap. Input files are distributed with the software.

**Figure 7.**
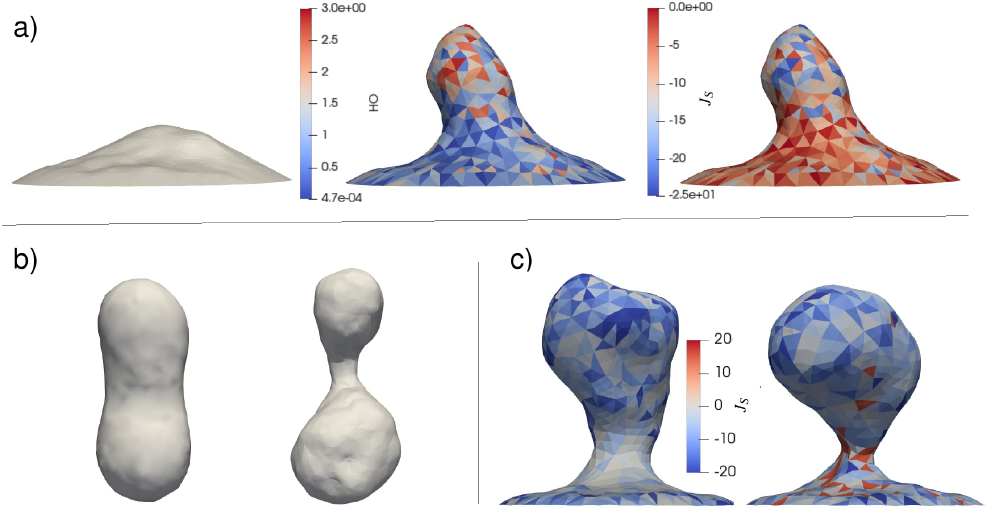
a) Coupling of lipid mixing and curvature generation with membrane geometry. Left panel: Simulation of lipid mixing without curvature, Central panel distribution of curved lipid and budding triggered by the same. Right panel, spontaneous curvature. b) ADE simulation starting from prolate geometry (left), as implemented by a global curvature restraint. c) Comparison of lipid budding and constriction with high spontaneous curvature (left), and neck-constricting protein mimicks (right)

Apart from a restriction on volume and surface integrals, it is also possible to restrict the mean curvature integral. This is equivalent to the ADE formulation, as area differences create global curvature restrictions. One example of such an ADE simulation is shown in Fig. 7 b). The left side is the starting geometry and only the curvature restriction enforces the panhandle shape. We have also obtained all manners of triple-panhandle shapes. Finally, it is possible to slightly relax the penalties to bring forth more flexibility. Under these conditions, it is possible to create and constrict multiple buds from a small spherical cap and generate deformations on buddings, etc. One example is shown in Fig. 7 c). This example points towards the possibilities that can be reached by the refinement of moves and the introduction of explicit proteins.

## CONCLUSION

We describe a method for the mesoscopic modeling of lipid membranes and related systems, based on a powerful but simple discretization scheme. This scheme is introduced here to address previous weaknesses of the Nagata interpolant and provides a new approach to DTS simulations. We also present a feature-rich implementation in the form of a Monte Carlo code. This software is capable of finding the well-known rotational solutions of the Helfrich energy and empirically exhibits the correct limits of stability. In its present form, the method is potentially useful for predicting e.g. the effect of lipid compositions, boundaries and proteins in a number of stylized ways. Our software, OrganL, can be used to study viral budding, curvature sorting and cellular shapes, membrane shape fluctuations and many other effects. The implementation design allows for easy extension of the code to include new couplings, adding e.g. a cytoskeleton. Current limitations are the lack of explicit surface-surface interactions beyond collisions (such as protein membrane interactions), explicit forces for fluid-structure interaction and topological changes, such as membrane fusion.

## Supporting information

Supplementary Matetials

## AUTHOR CONTRIBUTIONS

CA designed the research and wrote the code. BF contributed to the collision detection. CA and MD carried out all simulations and analyzed the data. CA and MD wrote the article, with input and edits from BF.

## ACKNOWLEDGMENTS

CA and MD thank Charles University for support via the PRIMUS research project PRIMUS/20/SCI/015. The authors also thank Vojtěch Kubáč for his contributions regarding the visualization and informative early work, Petr Pelech and Hina Arif for proofreading.

## SUPPLEMENTARY MATERIAL

The Supplementary Material contains a detailed derivation of the interpolants, algorithmic details about collision detection, quadrature and parallel execution as well as data structures and a description of additional energy functionals. Until acceptance of the final manuscript the program, supplementary videos and manual are available from the corresponding author.

## Notes

### Competing Interest Statement

The authors have declared no competing interest.

