## Supplementary Matetials for "OrganL: Dynamic Triangulation of Biomembranes using Curved Elements"

### Supplementary Material for OrganL: OrganL: Dynamic Triangulation of Biomembranes using Curved Elements

##### Edge Interpolant

The conditions given in the paper can be reformulated as a linear system of equations for the unknown coefficients $\boldsymbol{c}_1$ and $\boldsymbol{c}_2$ as follows

$$\mathbb{M}\boldsymbol{c} = \begin{bmatrix} \boldsymbol{n}_A \cdot \boldsymbol{d} \\ -\boldsymbol{n}_B \cdot \boldsymbol{d} \\ 0 \\ 0 \end{bmatrix}, \tag{1}$$

where $\boldsymbol{c} \overset{\text{def}}{=} [c_1^{(1)}, c_1^{(2)}, c_1^{(3)}, c_2^{(1)}, c_2^{(2)}, c_2^{(3)}]^\top$ and

$$\mathbb{M} \overset{\text{def}}{=} \begin{bmatrix} \boldsymbol{n}_A^\top, \boldsymbol{n}_A^\top \\ \boldsymbol{n}_B^\top, 2\boldsymbol{n}_B^\top \\ (\boldsymbol{d} \times \boldsymbol{n}_A)^\top, (\boldsymbol{d} \times \boldsymbol{n}_A)^\top \\ (\boldsymbol{d} \times \boldsymbol{n}_B)^\top, 2\,(\boldsymbol{d} \times \boldsymbol{n}_B)^\top \end{bmatrix}. \tag{2}$$

Explicitly, the pseudoinverse $\mathbb{M}^+$ determining the cubic interpolant is therefore

$$\mathbb{M}^+ = \lim_{\alpha \to 0^+} \begin{bmatrix} \boldsymbol{n}_A & \boldsymbol{n}_B & \boldsymbol{d} \times \boldsymbol{n}_A & \boldsymbol{d} \times \boldsymbol{n}_B \\ \boldsymbol{n}_A & 2\boldsymbol{n}_B & \boldsymbol{d} \times \boldsymbol{n}_A & 2\boldsymbol{d} \times \boldsymbol{n}_B \end{bmatrix}$$
$$\left( \begin{bmatrix} \boldsymbol{n}_A^\top, \boldsymbol{n}_A^\top \\ \boldsymbol{n}_B^\top, 2\boldsymbol{n}_B^\top \\ (\boldsymbol{d} \times \boldsymbol{n}_A)^\top, (\boldsymbol{d} \times \boldsymbol{n}_A)^\top \\ (\boldsymbol{d} \times \boldsymbol{n}_B)^\top, 2\,(\boldsymbol{d} \times \boldsymbol{n}_B)^\top \end{bmatrix} \begin{bmatrix} \boldsymbol{n}_A & \boldsymbol{n}_B & \boldsymbol{d} \times \boldsymbol{n}_A & \boldsymbol{d} \times \boldsymbol{n}_B \\ \boldsymbol{n}_A & 2\boldsymbol{n}_B & \boldsymbol{d} \times \boldsymbol{n}_A & 2\boldsymbol{d} \times \boldsymbol{n}_B \end{bmatrix} + \alpha \mathbb{I} \right)^{-1}. \tag{3}$$

The matrix inverse inside the brackets can be obtained analytically. For clarity and later numerical use, we introduce the notation

$$a \stackrel{\text{def}}{=} \boldsymbol{n}_A \cdot \boldsymbol{n}_B, \tag{4a}$$
$$b \stackrel{\text{def}}{=} \boldsymbol{n}_A \cdot (\boldsymbol{d} \times \boldsymbol{n}_B), \tag{4b}$$
$$c \stackrel{\text{def}}{=} \|\boldsymbol{d} \times \boldsymbol{n}_A\|^2, \tag{4c}$$
$$d \stackrel{\text{def}}{=} (\boldsymbol{d} \times \boldsymbol{n}_A) \cdot (\boldsymbol{d} \times \boldsymbol{n}_B), \tag{4d}$$
$$e \stackrel{\text{def}}{=} \|\boldsymbol{d} \times \boldsymbol{n}_B\|^2. \tag{4e}$$

Inserting these definitions the pseudoinverse is given by

$$\begin{aligned}\mathbb{M}^+ &= \lim_{\alpha\to 0^+} \begin{bmatrix} \boldsymbol{n}_A & \boldsymbol{n}_B & \boldsymbol{d}\times\boldsymbol{n}_A & \boldsymbol{d}\times\boldsymbol{n}_B \\ \boldsymbol{n}_A & 2\boldsymbol{n}_B & \boldsymbol{d}\times\boldsymbol{n}_A & 2\boldsymbol{d}\times\boldsymbol{n}_B \end{bmatrix} \left( \begin{bmatrix} 2 & 3a & 0 & 3b \\ 3a & 5 & -3b & 0 \\ 0 & -3b & 2c & 3d \\ 3b & 0 & 3d & 5e \end{bmatrix} + \alpha\mathbb{I} \right)^{-1} \\ &= \begin{bmatrix} \boldsymbol{n}_A & \boldsymbol{n}_B & \boldsymbol{d}\times\boldsymbol{n}_A & \boldsymbol{d}\times\boldsymbol{n}_B \\ \boldsymbol{n}_A & 2\boldsymbol{n}_B & \boldsymbol{d}\times\boldsymbol{n}_A & 2\boldsymbol{d}\times\boldsymbol{n}_B \end{bmatrix} \frac{1}{k}\mathbb{S},\end{aligned} \tag{5}$$

where $k \stackrel{\text{def}}{=} 81b^4 - 18(5c - 9ad + 5e)b^2 + \left(9a^2 - 10\right)\left(9d^2 - 10ce\right)$. This allows us to obtain an explicit expression for $\boldsymbol{c}$:

$$\boldsymbol{c} = \begin{bmatrix} \boldsymbol{n}_A & \boldsymbol{n}_B & \boldsymbol{d}\times\boldsymbol{n}_A & \boldsymbol{d}\times\boldsymbol{n}_B \\ \boldsymbol{n}_A & 2\boldsymbol{n}_B & \boldsymbol{d}\times\boldsymbol{n}_A & 2\boldsymbol{d}\times\boldsymbol{n}_B \end{bmatrix} \frac{1}{k}\mathbb{S} \begin{bmatrix} \boldsymbol{n}_A\cdot\boldsymbol{d} \\ -\boldsymbol{n}_B\cdot\boldsymbol{d} \\ 0 \\ 0 \end{bmatrix}. \tag{6}$$

The symmetric matrix $\mathbb{S}$ is defined as

$$\begin{bmatrix} -\left(5\left(9eb^2 + 9d^2 - 10ce\right)\right) & 27d\left(b^2 + ad\right) - 30ace & 45b(d - ae) & 3b\left(9b^2 - 10c + 9ad\right) \\ 27d\left(b^2 + ad\right) - 30ace & 20ce - 18\left(cb^2 + d^2\right) & -\left(3b\left(9b^2 + 9ad - 10e\right)\right) & 18b(ac - d) \\ 45b(d - ae) & -\left(3b\left(9b^2 + 9ad - 10e\right)\right) & -\left(5\left(9b^2 + \left(9a^2 - 10\right)e\right)\right) & 27a\left(b^2 + ad\right) - 30d \\ 3b\left(9b^2 - 10c + 9ad\right) & 18b(ac - d) & 27a\left(b^2 + ad\right) - 30d & -\left(2\left(9b^2 + \left(9a^2 - 10\right)c\right)\right), \end{bmatrix} \tag{7}$$

and can be comfortably evaluated without iterations.

#### Patch Interpolant

The three coordinates and three coefficients of the edge interpolants uniquely determine the patch interpolant. In order to use the cubic extension we propose here, we will follow the original approach as far as possible. The cubic surface interpolant

$$\boldsymbol{x}(\eta,\zeta) = \widetilde{\boldsymbol{c}_{00}} + \widetilde{\boldsymbol{c}_{10}}\eta + \widetilde{\boldsymbol{c}_{01}}\zeta + \widetilde{\boldsymbol{c}_{11}}\eta\zeta + \widetilde{\boldsymbol{c}_{20}}\eta^2 + \widetilde{\boldsymbol{c}_{02}}\zeta^2 + \widetilde{\boldsymbol{c}_{21}}\eta^2\zeta + \widetilde{\boldsymbol{c}_{12}}\eta\zeta^2 + \widetilde{\boldsymbol{c}_{30}}\eta^3 + \widetilde{\boldsymbol{c}_{03}}\zeta^3 \tag{8}$$

is the natural extension of the quadratic patch, again we require the interpolants at the edges

$$\boldsymbol{x}(\eta,0) = \boldsymbol{x}_{00} + (\boldsymbol{d}_1 - \boldsymbol{c}_{11} - \boldsymbol{c}_{12})\,\eta + \boldsymbol{c}_{11}\eta^2 + \boldsymbol{c}_{12}\eta^3, \tag{9a}$$
$$\boldsymbol{x}(1,\zeta) = \boldsymbol{x}_{10} + (\boldsymbol{d}_2 - \boldsymbol{c}_{21} - \boldsymbol{c}_{22})\,\zeta + \boldsymbol{c}_{21}\zeta^2 + \boldsymbol{c}_{22}\zeta^3, \tag{9b}$$
$$\boldsymbol{x}(\eta,\eta) = \boldsymbol{x}_{00} + (\boldsymbol{d}_3 - \boldsymbol{c}_{31} - \boldsymbol{c}_{32})\,\eta + \boldsymbol{c}_{31}\eta^2 + \boldsymbol{c}_{32}\eta^3, \tag{9c}$$

to coincide with the interpolant (8) restricted to the edges

$$\boldsymbol{x}(\eta,0) = \widetilde{\boldsymbol{c}_{00}} + \widetilde{\boldsymbol{c}_{10}}\eta + \widetilde{\boldsymbol{c}_{20}}\eta^2 + \widetilde{\boldsymbol{c}_{30}}\eta^3, \tag{10a}$$

$$\boldsymbol{x}(1,\zeta) = (\widetilde{\boldsymbol{c}_{00}} + \widetilde{\boldsymbol{c}_{10}} + \widetilde{\boldsymbol{c}_{20}} + \widetilde{\boldsymbol{c}_{30}}) + (\widetilde{\boldsymbol{c}_{01}} + \widetilde{\boldsymbol{c}_{11}} + \widetilde{\boldsymbol{c}_{21}})\,\zeta + (\widetilde{\boldsymbol{c}_{02}} + \widetilde{\boldsymbol{c}_{12}})\,\zeta^2 + \widetilde{\boldsymbol{c}_{03}}\zeta^3, \tag{10b}$$

$$\boldsymbol{x}(\eta,\eta) = \widetilde{\boldsymbol{c}_{00}} + (\widetilde{\boldsymbol{c}_{10}} + \widetilde{\boldsymbol{c}_{01}})\,\eta + (\widetilde{\boldsymbol{c}_{11}} + \widetilde{\boldsymbol{c}_{20}} + \widetilde{\boldsymbol{c}_{02}})\,\eta^2 + (\widetilde{\boldsymbol{c}_{21}} + \widetilde{\boldsymbol{c}_{12}} + \widetilde{\boldsymbol{c}_{30}} + \widetilde{\boldsymbol{c}_{03}})\,\eta^3. \tag{10c}$$

By equating the corresponding coefficients we arrive at

$$\widetilde{\boldsymbol{c}_{00}} = \boldsymbol{x}_{00}, \tag{11a}$$
$$\widetilde{\boldsymbol{c}_{10}} = \boldsymbol{d}_1 - \boldsymbol{c}_{11} - \boldsymbol{c}_{12}, \tag{11b}$$
$$\widetilde{\boldsymbol{c}_{20}} = \boldsymbol{c}_{11}, \tag{11c}$$
$$\widetilde{\boldsymbol{c}_{30}} = \boldsymbol{c}_{12}, \tag{11d}$$
$$\widetilde{\boldsymbol{c}_{00}} + \widetilde{\boldsymbol{c}_{10}} + \widetilde{\boldsymbol{c}_{20}} + \widetilde{\boldsymbol{c}_{30}} = \boldsymbol{x}_{10}, \tag{11e}$$
$$\widetilde{\boldsymbol{c}_{01}} + \widetilde{\boldsymbol{c}_{11}} + \widetilde{\boldsymbol{c}_{21}} = \boldsymbol{d}_2 - \boldsymbol{c}_{21} - \boldsymbol{c}_{22}, \tag{11f}$$
$$\widetilde{\boldsymbol{c}_{02}} + \widetilde{\boldsymbol{c}_{12}} = \boldsymbol{c}_{21}, \tag{11g}$$
$$\widetilde{\boldsymbol{c}_{03}} = \boldsymbol{c}_{22}, \tag{11h}$$
$$\widetilde{\boldsymbol{c}_{00}} = \boldsymbol{x}_{00}, \tag{11i}$$
$$\widetilde{\boldsymbol{c}_{10}} + \widetilde{\boldsymbol{c}_{01}} = \boldsymbol{d}_3 - \boldsymbol{c}_{31} - \boldsymbol{c}_{32}, \tag{11j}$$
$$\widetilde{\boldsymbol{c}_{11}} + \widetilde{\boldsymbol{c}_{20}} + \widetilde{\boldsymbol{c}_{02}} = \boldsymbol{c}_{31}, \tag{11k}$$
$$\widetilde{\boldsymbol{c}_{21}} + \widetilde{\boldsymbol{c}_{12}} + \widetilde{\boldsymbol{c}_{30}} + \widetilde{\boldsymbol{c}_{03}} = \boldsymbol{c}_{32}. \tag{11l}$$

Equations (11a)–(11d) and (11h) immediately determine the coefficients $\widetilde{\boldsymbol{c}_{00}}$, $\widetilde{\boldsymbol{c}_{10}}$, $\widetilde{\boldsymbol{c}_{20}}$, $\widetilde{\boldsymbol{c}_{30}}$, and $\widetilde{\boldsymbol{c}_{03}}$. Equation (11e) is just a sum of the first four equations so we can forget about this equation as well as about equation (11i). Equation (11j) finally determines the coefficient $\widetilde{\boldsymbol{c}_{01}}$ so that we obtain

$$\widetilde{\boldsymbol{c}_{00}} = \boldsymbol{x}_{00}, \tag{12a}$$
$$\widetilde{\boldsymbol{c}_{10}} = \boldsymbol{d}_1 - \boldsymbol{c}_{11} - \boldsymbol{c}_{12}, \tag{12b}$$
$$\widetilde{\boldsymbol{c}_{01}} = \boldsymbol{d}_2 + \boldsymbol{c}_{11} + \boldsymbol{c}_{12} - \boldsymbol{c}_{31} - \boldsymbol{c}_{32} \tag{12c}$$
$$\widetilde{\boldsymbol{c}_{20}} = \boldsymbol{c}_{11}, \tag{12d}$$
$$\widetilde{\boldsymbol{c}_{30}} = \boldsymbol{c}_{12}, \tag{12e}$$
$$\widetilde{\boldsymbol{c}_{03}} = \boldsymbol{c}_{22}, \tag{12f}$$

Plugging the known coefficients into equations (11f), (11g), (11k), and (11l) we arrive at a system of 4 equations

$$\widetilde{\boldsymbol{c}_{11}} + \widetilde{\boldsymbol{c}_{21}} = \boldsymbol{c}_{31} + \boldsymbol{c}_{32} - \boldsymbol{c}_{11} - \boldsymbol{c}_{12} - \boldsymbol{c}_{21} - \boldsymbol{c}_{22}, \tag{13a}$$
$$\widetilde{\boldsymbol{c}_{02}} + \widetilde{\boldsymbol{c}_{12}} = \boldsymbol{c}_{21}, \tag{13b}$$
$$\widetilde{\boldsymbol{c}_{11}} + \widetilde{\boldsymbol{c}_{02}} = \boldsymbol{c}_{31} - \boldsymbol{c}_{11}, \tag{13c}$$
$$\widetilde{\boldsymbol{c}_{21}} + \widetilde{\boldsymbol{c}_{12}} = \boldsymbol{c}_{32} - \boldsymbol{c}_{12} - \boldsymbol{c}_{22} \tag{13d}$$

for the unknown coefficients $\widetilde{\boldsymbol{c}_{11}}$, $\widetilde{\boldsymbol{c}_{02}}$, $\widetilde{\boldsymbol{c}_{21}}$, and $\widetilde{\boldsymbol{c}_{12}}$. However, by summing equations (13c) and (13d) and subtracting Eq. (13b) we obtain (13a). As a result, the system is underdetermined, and the general solution is given by

$$\widetilde{\boldsymbol{c}_{11}} = \boldsymbol{c}_{31} - \boldsymbol{c}_{11} - \boldsymbol{c}_{21} + \xi, \tag{14a}$$
$$\widetilde{\boldsymbol{c}_{02}} = \boldsymbol{c}_{21} - \xi, \tag{14b}$$
$$\widetilde{\boldsymbol{c}_{21}} = \boldsymbol{c}_{32} - \boldsymbol{c}_{12} - \boldsymbol{c}_{22} - \xi, \tag{14c}$$
$$\widetilde{\boldsymbol{c}_{12}} = \xi, \tag{14d}$$

where $\xi \in \mathbb{R}$ parametrizes the individual solutions. To obtain a unique patch interpolant, we require that if the edge interpolants are all quadratic, i.e.

$$\boldsymbol{c}_{12} = \boldsymbol{c}_{22} = \boldsymbol{c}_{32} = \boldsymbol{0}, \tag{15}$$

then, the patch interpolant should be quadratic as well, i.e.

$$\widetilde{\boldsymbol{c}_{21}} = \widetilde{\boldsymbol{c}_{12}} = \widetilde{\boldsymbol{c}_{30}} = \widetilde{\boldsymbol{c}_{03}} = \boldsymbol{0}. \tag{16}$$

We see that if (15) holds, then $\widetilde{\boldsymbol{c}_{30}}$ and $\widetilde{\boldsymbol{c}_{03}}$ trivially vanish by virtue of Eq. (12e) and Eq. (12f). Further, under the assumption that (15) holds, see Eq. (14c) and Eq. (14d), reduce to

$$\widetilde{\boldsymbol{c}_{21}} = -\xi, \tag{17a}$$
$$\widetilde{\boldsymbol{c}_{12}} = \xi, \tag{17b}$$

and we immediately see that these coefficients vanish if and only if $\xi = 0$. Consequently, the unique cubic patch interpolant which reduces to quadratic patch interpolant if the edge interpolants are quadratic is obtained by setting $\xi = 0$. The final set of explicit formulae for the coefficients of Eq. (8) thus reads

$$\widetilde{\boldsymbol{c}_{00}} = \boldsymbol{x}_{00}, \tag{18a}$$
$$\widetilde{\boldsymbol{c}_{10}} = \boldsymbol{d}_1 - \boldsymbol{c}_{11} - \boldsymbol{c}_{12}, \tag{18b}$$
$$\widetilde{\boldsymbol{c}_{01}} = \boldsymbol{d}_2 + \boldsymbol{c}_{11} + \boldsymbol{c}_{12} - \boldsymbol{c}_{31} - \boldsymbol{c}_{32} \tag{18c}$$
$$\widetilde{\boldsymbol{c}_{20}} = \boldsymbol{c}_{11}, \tag{18d}$$
$$\widetilde{\boldsymbol{c}_{30}} = \boldsymbol{c}_{12}, \tag{18e}$$
$$\widetilde{\boldsymbol{c}_{03}} = \boldsymbol{c}_{22}, \tag{18f}$$
$$\widetilde{\boldsymbol{c}_{11}} = \boldsymbol{c}_{31} - \boldsymbol{c}_{11} - \boldsymbol{c}_{21}, \tag{18g}$$
$$\widetilde{\boldsymbol{c}_{02}} = \boldsymbol{c}_{21}, \tag{18h}$$
$$\widetilde{\boldsymbol{c}_{21}} = \boldsymbol{c}_{32} - \boldsymbol{c}_{12} - \boldsymbol{c}_{22}, \tag{18i}$$
$$\widetilde{\boldsymbol{c}_{12}} = \boldsymbol{0}. \tag{18j}$$

Thus, the patch interpolant is trivially and uniquely determined from the edge interpolants, with a natural fallback to the quadratic case. Such a natural integration enables us to use the cubic interpolants only where the normal arrangement demands it.

##### Collision Detection

Discounting co-planar triangles, an initial requirement for triangle-triangle intersection is that one vertex of triangle $T_1$ should be on the opposite side of the plane of $T_2$ (denoted as $\pi_2$) than the other two. The plane $\pi_2$ is given by the equations

$$N_2 \cdot X + d_2 = 0, \tag{19}$$

$$N_2 = (V_1^2 - V_0^2) \times (V_2^2 - V_0^2), \tag{20}$$

$$d_2 = -N_2 \cdot V_0^2, \tag{21}$$

where $V_i^n$ denotes the position of the $i$-th vertex of triangle $T_n$. The signed distances of $V_i^1$ to $\pi_2$ are given (up to a multiplicative factor) by

$$d_{V_i^1} = N_2 \cdot V_i^1 + d_2. \tag{22}$$

In case all $d_{V_i^1}$ have the same sign, then $T_1$ lies wholly on one side of $\pi_2$, and their overlap is ejected. The test must also be repeated for $T_2$ and $\pi_1$.

If the intersection can not be rejected based on the above two tests, the planes of the two triangles must intersect at a line $L = O + tD$, where $D = N_1 \times N_2$ is the direction of the line and $O$ is a point on it. Then, the two triangles intersect each other if their intersections with the line $L$ $\left[t_0^1, t_1^1\right]$ and $\left[t_0^2, t_1^2\right]$ overlap. The end points of the intervals are given by

$$t_0^1 = p_{V_0^1} + \left(p_{V_1^1} - p_{V_0^1}\right) \frac{d_{V_0^1}}{d_{V_0^1} - d_{V_1^1}}, \tag{23}$$

$$t_1^1 = p_{V_2^1} + \left(p_{V_2^1} - p_{V_0^1}\right) \frac{d_{V_0^1}}{d_{V_0^1} - d_{V_2^1}}, \tag{24}$$

$$t_0^2 = p_{V_0^2} + \left(p_{V_1^2} - p_{V_0^2}\right) \frac{d_{V_0^2}}{d_{V_0^2} - d_{V_1^2}}, \tag{25}$$

$$t_1^2 = p_{V_2^2} + \left(p_{V_2^2} - p_{V_0^2}\right) \frac{d_{V_0^2}}{d_{V_0^2} - d_{V_2^2}}, \tag{26}$$

where $p_{V_i^n} = D \cdot (V_i^n - O)$ is the projection of the $V_i^n$ triangle vertex onto the line $L$. Finally, in the rare case of co-planar, triangles a simple two-dimensional triangle-triangle intersection test is performed (see [1]).

Because we are not aware of existing methods to compute intersections between Nagata surfaces, we employ a subdivision scheme to refine the triangle faces into sub-faces. The sub-faces are created by introducing one additional point inside the middle of the triangle. Then, we detect the collision between the sub-faces of the triangles.

##### Geometric Cutoffs

There is an additional constraint to the move, which considers the face normals between the triangles modified by the vertex move and their neighbors. The zone affected by this is given by the blue triangles in the central panel of Fig. 4 of the main manuscript. We compute the face normals of these triangles by

$$\boldsymbol{n}_F = (\boldsymbol{x}_0 \boldsymbol{x}_1) \times (\boldsymbol{x}_0 - \boldsymbol{x}_2). \tag{27}$$

We ensure that these normals align with the orientation of the patch, create unit normals and then require these to fulfill

$$\hat{\boldsymbol{n}}_{F,i} \cdot \hat{\boldsymbol{n}}_{F,j} < \cos(\pi/6). \tag{28}$$

for each pair of faces $j$ adjacent to the vertex of an upgraded triangle $i$. This restriction serves to avoid the generation of singular edges, i.e. edges where the direction of normal vectors across the patch edge exhibits a strong discontinuity. We will address this issue in a more direct manner, in the penalty section. In addition, collision checks are performed.

The move shown in Fig. 4 c) of the main manuscript is the so-called "deep vertex move". This move constitutes a vertex move, followed by normal reconstruction. In the first part a trial vertex $\boldsymbol{v}_P$ is generated in the same way as for an ordinary vertex move. In the next step, the normals of all the triangles that contain the vertex $\boldsymbol{v}_P$ are updated via a direct estimation of the normal. The face normals adjacent to a vertex $\boldsymbol{x}_0$ are computed according to Eq. 27, normalized and weighted

$$\boldsymbol{n} = \sum_{i=1}^{2} \omega_i \hat{\boldsymbol{n}}_{F,i}; \qquad \boldsymbol{n}_P = \frac{\boldsymbol{n}}{\|\boldsymbol{n}\|}. \tag{29}$$

Several normal estimation algorithms are possible and differ by their choice of $\omega_i$. There is no consensus in the literature about this distinction.[2] All of these simple algorithms have linear convergence. We choose the algorithm proposed by Max[3]

$$\omega_i = \frac{\sin\alpha}{\|\boldsymbol{d}_1\|\|\boldsymbol{d}_2\|}. \tag{30}$$

Where the vectors $\boldsymbol{d}_i$ are the edge lengths. This was observed by Max to deliver better performance in unequally spaced meshes. The validity checks for normal moves and vertex moves for the newly generated geometry are then applied. Note that, as the normal vectors around the moved vertex are updated, triangles incorporating these normals have to be updated as well, which gives the move a higher "depth" in the mesh.

#### Energy Evaluation

The surface integrals $\Phi$ of any function $f$ over a patch interpolant $\boldsymbol{x}$ are

$$\Phi = \int_0^1 d\eta \int_0^\eta d\zeta f(\eta,\zeta)\|\boldsymbol{x}_\eta(\eta,\zeta) \times \boldsymbol{x}_\zeta(\eta,\zeta)\|. \tag{31}$$

They be summed up to yield an integral over the entire surface. Unfortunately, it is not generally possible to analytically evaluate $\Phi$. The integral is therefore discretized on the domain of the interpolant:

$$\Phi \approx \sum_i \omega_i f(\boldsymbol{p}_i)\|\boldsymbol{x}_\eta(\boldsymbol{p}_i) \times \boldsymbol{x}_\zeta(\boldsymbol{p}_i)\|. \tag{32}$$

The integration is performed via Gaussian quadrature. The points $\boldsymbol{p}_i = [\zeta_i, \eta_i]$ and the weights $\omega_i$ are taken from the the literature.[4] In particular, the quadrature weights are chosen to lie away from the edges and are symmetrical on the domain. Special care is given to the evaluation of the evaluation of $H$ and $K$ on the surface. These quantities can be obtained from the Weingarten equations, yielding

$$H(\eta,\zeta) = \frac{LG - MF + NE}{2(EG - F^2)}, \tag{33}$$

and

$$K_G(\eta,\zeta) = \frac{LN - M^2}{EG - F^2}. \tag{34}$$

Here, $E, F, G$ are the matrix elements of the first fundamental form, which is defined by $I(i,j) = \boldsymbol{x}_i \cdot \boldsymbol{x}_j$ and $L, N, M$ are the elements of the second fundamental form, given by $II(i,j) = \boldsymbol{x}_{ij} \cdot \hat{\boldsymbol{n}}$. These coefficients can be trivially computed from $\boldsymbol{x}(\eta,\zeta)$. For the quadratic patch, the second derivatives are directly related to the coefficients of the edge interpolants.

$$\boldsymbol{x}_{\eta\eta} = \boldsymbol{c}_1, \tag{35a}$$

$$\boldsymbol{x}_{\zeta\zeta} = \boldsymbol{c}_2, \tag{35b}$$

$$\boldsymbol{x}_{\eta\zeta} = \boldsymbol{c}_3 - \boldsymbol{c}_1 - \boldsymbol{c}_2, \tag{35c}$$

For the cubic interpolant, the second derivatives will still depend on position. Further simplifications can be made for any closed surface, as by the Gauss-Bonnet theorem

$$\int_{Surf.} K_G = 2\pi\chi(M), \tag{36}$$

with $\chi(M)$ being the Euler characteristic. This characteristic is not changed unless the topology of the membrane is modified. Therefore, the corresponding term in the evaluation of $F$ can be neglected.

##### Penalties

The global volume and area are implemented via quadratic penalties, with constants $\lambda$:

$$E_{Pen.} = \frac{\lambda_0}{2}\left(A_0 - \int_{Surf.} \mathrm{d}A\right)^2 + \frac{\lambda_1}{2}\left(V_0 - \int_{Vol} \mathrm{d}V\right)^2 \tag{37}$$

This can be interpreted as letting the surface tension $\sigma$ is computed from an area compressibility modulus $K_A = \lambda_0$

$$\sigma = K_A(A - A_0), \tag{38}$$

to increase realism. Similarly, the pressure term includes a compressibility term $\kappa_T = \lambda_1$

$$p = -\kappa_T(V - V_0). \tag{39}$$

Turning the Lagrange multipliers into penalty functions facilitates implementation while at the same time increasing the realism of the physical model.

##### Scheduling and Parallelization

The short range of interpolation facilitates the parallelization of our code. Our long-term goal for OrganL is the simulation of entire organelles, which contain numerous processes coupled to the membrane geometry. In particular, lipid mixing, the cytoskeleton and membrane proteins and long-range interactions, such as electrostatics should therefore to some degree be anticipated by the code. For example, it would make sense, to e.g. also mesh membrane proteins and other structures with the Nagata interpolation, but restrict their flexibility compared to that of lipid membranes or add additional energy terms, such as area difference elasticity or nonideal mixing, see Fig. 1 to see how these thoughts are reflected in our implementation. The challenge in this context is to properly encapsulate the code, so that these logical extensions do not need extensive rewrites in different parts of the code. In particular, parallelization should be facilitated by the structure of the program. The first requirement for efficient parallelization is the generation of neighbor lists, which enable the localization of through-space interactions.

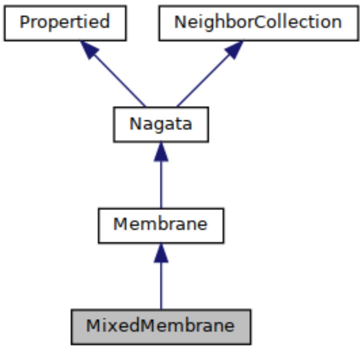


Figure 1: Class inheritance diagram for a Membrane with lipid Mixing, where the *Nagata* object incorporates the interfaces necessary for simulation moves in an abstract fashion.

On the other hand, it is convenient to have access to an automatic update mechanism for dependent objects, answering questions of the type "What objects do need to be updated, if a vertex is modified". Examples of such dependent objects could be transmembrane channels, or parts of the cytoskeleton, which are directly coupled to membrane structure, but may not possess significant through-space interactions, as well as coupled monolayers. Yet, these objects may also come with their own neighbors, and this can make it challenging to schedule the parallel execution of a move. Our paradigm is to apply Monte-Carlo moves to different *Nagata* entities, which have properties. These *Properties* in turn define the moves that can be applied to them. In Fig. 1 the *Propertied* and *NeighborCollection* classes provide interfaces to automatic updates and neighbor lists. In Fig. 2, an overview is given on the different types of properties exported by a *Membrane* by default. The *PropertyMap* object allows to query and access these by type. Note, how the actual mesh as well as the way energy is evaluated on the mesh are encapsulated away. Both the energy functional and the constituent properties of the membrane can be modified at run-time, so that for most extensions this part of the codebase has to be neither touched nor subclassed.

To give an example, the individual *VertexProperty* objects provide direct pointers to the vertex as stored on the mesh. They also store the *Dependencies* of each Vertex in a list in the form of pointers to *Properties*. The basic dependencies here are the *Faces*, which contain the interpolant and are connected to each vertex, but other dependencies could be added at run time. Each property has an update trigger, that recursively updates all the *Dependencies*. Finally, each property indicates, whether it contains a contribution to the energy or is subject

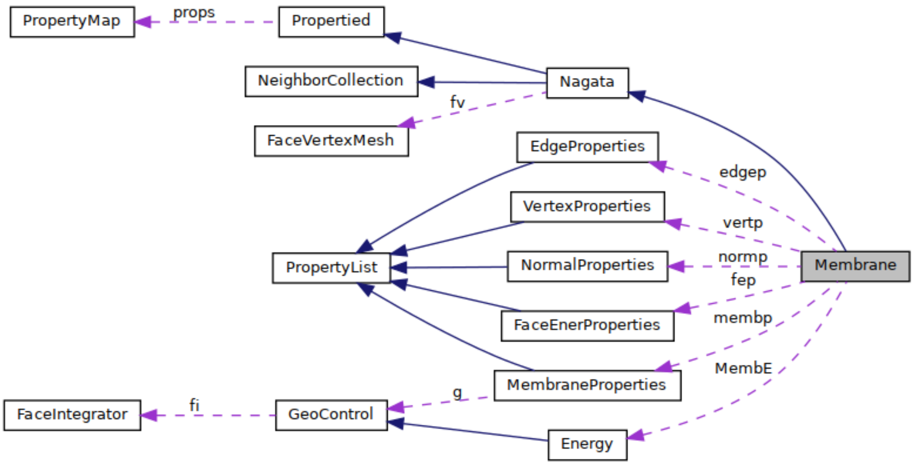


Figure 2: Class inheritance diagram for a Membrane with lipid Mixing, where the *Nagata* object incorporates the interfaces necessary for simulation moves in an abstract fashion.

to collisions. Searching the dependency tree of a *Property* allows to identify all objects which might cause trouble when being updated by concurrent moves. Avoiding race conditions demands that moves that modify the same objects cannot be executed at the same time. Using the aforementioned data structures, we define a general parallel Monte-Carlo move scheduling in the *MCMove* class, which gains access to the *Nagata* object via a *Simulation* object. The latter also stores various run parameters, such as the total number of steps and the sequence of moves to be executed. The Monte-Carlo moves are scheduled as follows

**procedure** CREATEPARALLELPLAN(Simulation $s$)
  **for all** Nagata $n \in s$ **do**
    **if** $n$ has set $A$ of compatible properties $P$ **then**
      Array $R[Rounds][Indices] \leftarrow \varnothing$
      Array $G[Indices][Indices] \leftarrow \varnothing$
      $r \leftarrow 0$
      **while** $A \neq \varnothing$ **do**
        $t \leftarrow 0$
        $l \leftarrow$Sizeof($P$)
        $D \leftarrow \varnothing$
        Array $M[Dim = l] \leftarrow \varnothing$
        **while** $t \leq T_{Max}$ **do**
          $p \leftarrow$Random $\in A$
          $d \leftarrow$EnergeticDependencies($p$)
          **if** $d \bigcap D = \varnothing$ **then**
            $D \leftarrow D \bigcup d$
            $R[r] \leftarrow R[r] \bigcup$index($p$)
            $G[p] \leftarrow d$
            $A \leftarrow\subset A \subsetneq index[p]$
          **end if**

```
                t ← t + 1
            end while
            r ← r + 1
        end while
    end if
end for
end procedure
```

At the end array $R$ contains a round $r$. Each round $r$ is a list of properties, whose moves can be performed simultaneously. The dependencies of each property are read out and stored in $G$. The reason for this is that while the property automatically updates, e.g. the underlying triangles, there is currently no such mechanism for energy retrieval and validation. Hence, $G$ is passed on together with $R$ for execution. It is obvious, that load balancing and rescheduling can be performed via changing the composition of $R$. However, in our case, we stay with one $R$ which is cycled via $r$ during move execution. Scheduling is also performed in parallel, with only assignments being taken out of the loop. The Simulation object then executes each move in the following way:

```
procedure ExecuteMoves(Nagata n, R, r, G, Neighbourlist b)
    for all p ∈ R[r] do
        repeat
            p_O ← p
            p ←RandomMove(p)
            RecursiveUpdate(p)
            ΔE ← n::ComputeEnergyChange(p, b, G[p])
            if CheckAccepted(ΔE) and CheckAllowed(p, G[p]) then
                if NoCollision(p, b) then
                    p ← Accepted
                    SaveEnergyChange(p, b)
                end if
            end if
            p ← p_O
            RecursiveUpdate(p)
        until p is accepted
    end for
```

This for loop is parallel. Timing issues can and do occur, with the different moves finishing at different times, however, in practice good scaling is achieved over 48 cores. The performance will obviously depend on the number of mesh points vs the number of cores etc. In practice, the computation of the energy change is done while querying the energy changes for the set $G[p]$ from the underlying Nagata object. The predicted energy change for each element of $G[p]$ is stored on the mesh and is moved to the final energy upon acceptance of the move. As Fig. 2 shows, the way energy is calculated is itself an object, that refers to the integrators. The change in energy may involve global restraints of the type encountered in the main text. These are also accessible as properties of the *Membrane*, e.g. the *MembraneProperties* allow for accessing globals, such as Volume, Area integrals as well as their defaults and penalties $\lambda$. So in order to compute the change in energy, these are required together with the $\Delta\Phi$ computed on the fly. The updating of these global properties together with the total energy is excluded from parallelization. Performing all the MC scheduling and energy changes through a Nagata object removes the need to modify the MC code when changing the energy functional, and provides a template for the implementation of more sophisticated MC moves.

##### Alternative Energy Functionals

This computer code has been written with the ease of implementation for new functionality in mind. In particular, we have already implemented several energy functionals, such as the pure Helfrich functional

$$F = \int_{Surf.} dA \left\{ \frac{\kappa}{2}(H - J_S)^2 \right\} \tag{40}$$

as well as quadratic double constraints

$$\int_{Surf.} dA \left\{ \frac{\kappa}{2}(H - J_S)^2 + \frac{K_A}{2}(A_i - A_0)^2 + \sigma \right\} + \int_{Vol} dVp. \tag{41}$$

In this case $K_A$ and $A_0$ are local and stored on the faces, and $A_i$ is computed for each face. Other functionals, such as the version of area difference elasticity[5]

$$\int_{Surf.} dA \left\{ \frac{\kappa}{2}(H - J_S)^2 + \lambda H + \sigma \right\} + \int_{Vol} dVp \tag{42}$$

have been also implemented in addition to the standard Helfrich energy as described in the main manuscript. Another quadratic penalty is added to the mean curvature integral as described above. Note, how the addition of a new global dependency is possible without subclassing the *Membrane* object. It is completely sufficient to overload the *Energy*, of a *Membrane* which can be done at run time. This opens the space for experimentation with new types of energies. Because the functionals are also able to choose their own integrators, it should easily be possible to further localize the parameters, for example by storing coefficients of functions on the faces. An important limit to consider is merely the minimally quadratic degree of the supporting membrane function.
